# Enzymes define pathways and metabolic relationships

**DOI:** 10.1101/2025.06.16.657199

**Authors:** Jiada Zhan, Zachery R. Jarrell, Melissa L. Kemp, Jaclyn Weinberg, Young-Mi Go, Greg S. Martin, Dean P. Jones

## Abstract

Gene-centric pathway mapping tools, widely used to interpret untargeted liquid chromatography mass spectrometry (LC-MS) metabolomics data, may underperform because a single metabolite can generate multiple spectral features, inflating false positive rates. Classic enzymology, which established metabolite flow before gene sequencing, offers experimentally validated precursor-product relationships that could overcome these ambiguities. We evaluated whether enzymology-defined precursor-product correlations are consistently detectable in human plasma LC-MS data. We detected amino acids, carnitine-related, TCA cycle, and pentose phosphate pathway metabolites in one individual sampled eight times over five years and in 50 adults sampled 6 to 8 times each. In the single participant repeated measures, strong positive correlations were observed for most direct precursor-product pairs. The longitudinal and cross-sectional analyses reproduced these patterns. Precursor-product proportionality, a fundamental principle of enzymology, is detectable in LC-MS datasets and remains consistent across studies. Applying these correlations to metabolomics workflows can improve pathway analysis, help metabolite identification, and reveal how genetic variations, diets, therapeutic drugs, and environmental exposures jointly impact metabolic pathways.

## Introduction

Recently, five biochemical and analytical challenges have been identified that can cause gene-centric tools to underperform when applied to metabolomics data.^1^ Untargeted liquid chromatography mass spectrometry (LC-MS) metabolomics-based pathway analysis often relies on pathway-mapping tools,^2,3^ but ambiguities in mass spectral peak assignment – multiple peaks arising from the same metabolite – can generate high false positive rates.^4,5^ Most metabolic pathways were established through enzymology before gene sequencing became available, and these studies offer direct, experimentally validated maps of metabolite flow. Thus, enzymology offers a simple but powerful way forward to enhance pathway analysis.

Enzymology established that metabolites flow from precursors to products and form dynamic linear, cyclical, and branched systems with regulation occurring at committed steps and branch points (**Fig. 1a)**.^6^ Here, we define a precursor-product relationship as a one-step biochemical conversion. Biochemical analyses showed that under steady-state conditions, the concentrations of metabolites often vary proportionately (see legend, **Fig. 1a)**.^6,7^ This principle was demonstrated in the pathway-level identification of xenobiotics and xenobiotic metabolites in blood^9^ and endogenous metabolites in plant tissue.^7^

**Figure 1.**
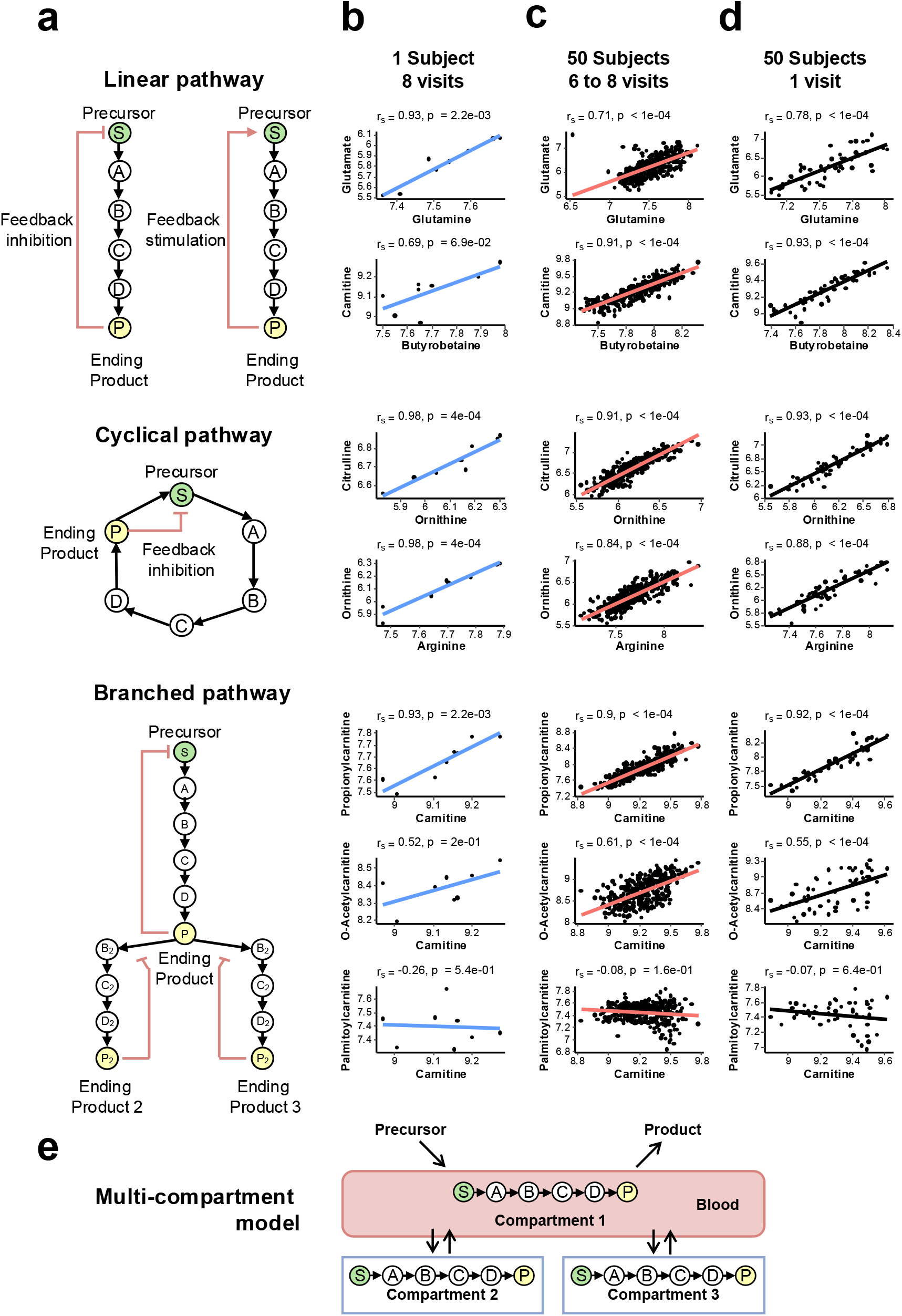
Precursor-product relationships in human plasma. (**a**). Metabolism occurs in linear (left, top), cyclic (left, middle), and branched pathways (left, bottom). For sequential enzyme reactions in an open system at steady state, rates of precursor (S) input, formation of intermediate products (A, B, C, D), and formation and elimination of end product (P) are equivalent. With the same reaction rates for all steps, steady-state concentrations of intermediates [A, B, C, D] are related by the respective pseudo-first order rate constants (k<> _j_): k_1_[S] = k_2_[A] = k_3_[B] = k_4_[C] = k_5_[D]. Because regulation typically occurs only at branch points or at sites committing a hydrocarbon to a specific fate, this means that proportional relationships of precursors and one-step intermediate products are expected. Noted, when there is no intermediate product between a precursor and end product, such as glutamine and glutamate, the notation of intermediate products should be disregarded. (**b**) Proportional intensities for precursor and product metabolites in different pathways are evident in eight repeated blood samplings from an individual collected over five years (blue). Examples are provided for plasma amino acids and carnitine-related metabolites, measured by hydrophilic interaction liquid chromatography (HILIC) and mass spectrometry (LC-MS) with positive ESI (See Reference 10 for methodological details)^24^. All the metabolites on the x-axis are the precursors, and all the metabolites on the y-axis are the products. Glutamine, glutamate, butyrobetaine, and carnitine are considered part of a linear pathway, ornithine, citrulline, and arginine are considered part of a cyclical pathway (urea cycle), and carnitine, propionylcarnitine, O-acetylcarnitine,and palmitoylcarnitine are considered part of a branched pathway (carnitine shuttle). Sample collection involved overnight fasting for the first blood draw, but not for the others. Note that an example of a precursor-product pair without proportional concentrations, palmitoylcarnitine and carnitine (bottom row), is provided. (**c**) Proportional intensities for the same precursor and product pairs as in Panel b for 50 individuals with multiple repeated blood samplings across five years (red). (**d**) Cross-sectional analysis of baseline samples for the 50 individuals in Panel **c** (black) shows that precursor-product correlations are preserved for the same precursor and product pairs. (**e**) In multicompartment models, most metabolism occurs in organ compartments rather than the blood. Despite this, the data in Panels **b** to **d** show that proportional relationships of precursors and products are observed in blood plasma samples.

## Results

Here, we show examples of precursor-product correlations observed from hydrophilic interaction liquid chromatography (HILIC) coupled to mass spectrometry with positive electrospray ionization (ESI) for amino acids (glutamine, glutamate, ornithine, citrulline, and arginine) and carnitine-related metabolites (butyrobetaine, carnitine, propionylcarnitine, O-acetylcarnitine,palmitoylcarnitine) in one person’s plasma sampled eight times across five years (**Fig. 1b**).^9^ These metabolites were confirmed by matching retention time (RT) and ion dissociation spectra (MS/MS) to authentic standards. Among these examples, the lack of proportional change of palmitoylcarnitine with carnitine illustrated that a positive correlation is not always present, especially for metabolites at branch points and regulated sites.

To test whether positive precursor-product correlations exist at a population level, we analyzed LC-MS plasma data from 50 individuals, each with six to eight visits.^9^ Consistent with the data found in the individual (**Fig. 1b**), amino acids and carnitine-related metabolites exhibit positive correlations in longitudinal analyses across the 50 individuals (**Fig. 1c**), except for the lack of proportional variation of palmitoylcarnitine with carnitine. Unlike these longitudinal comparisons, however, most human metabolomics studies use a cross-sectional design with only single LC-MS snapshots of individuals. To determine whether enzymology-defined precursor-product correlations are present in a cross-sectional sampling, we restricted comparisons of the 50 individuals to include only baseline samples, and results showed the same relationships were present (**Fig. 1d**).

To determine if these trends are upheld with reversed phase chromatography coupled to negative ESI, we performed additional analyses for the TCA cycle (isocitric acid and alpha-ketoglutaric acid) and pentose phosphate pathway metabolites (D-gluconate and 6-phosphogluconic acid) with more subjects included. Isocitric acid and alpha-ketoglutaric acid showed significant positive correlations in the longitudinal and cross-sectional analyses (**Fig. 2a**). A significant correlation was observed between 6-phosphogluconic acid and D-gluconate (**Fig. 2b**), but not between D-gluconate and D-glucono-1,5-lactone (**Fig. 2b**).

**Figure 2.**
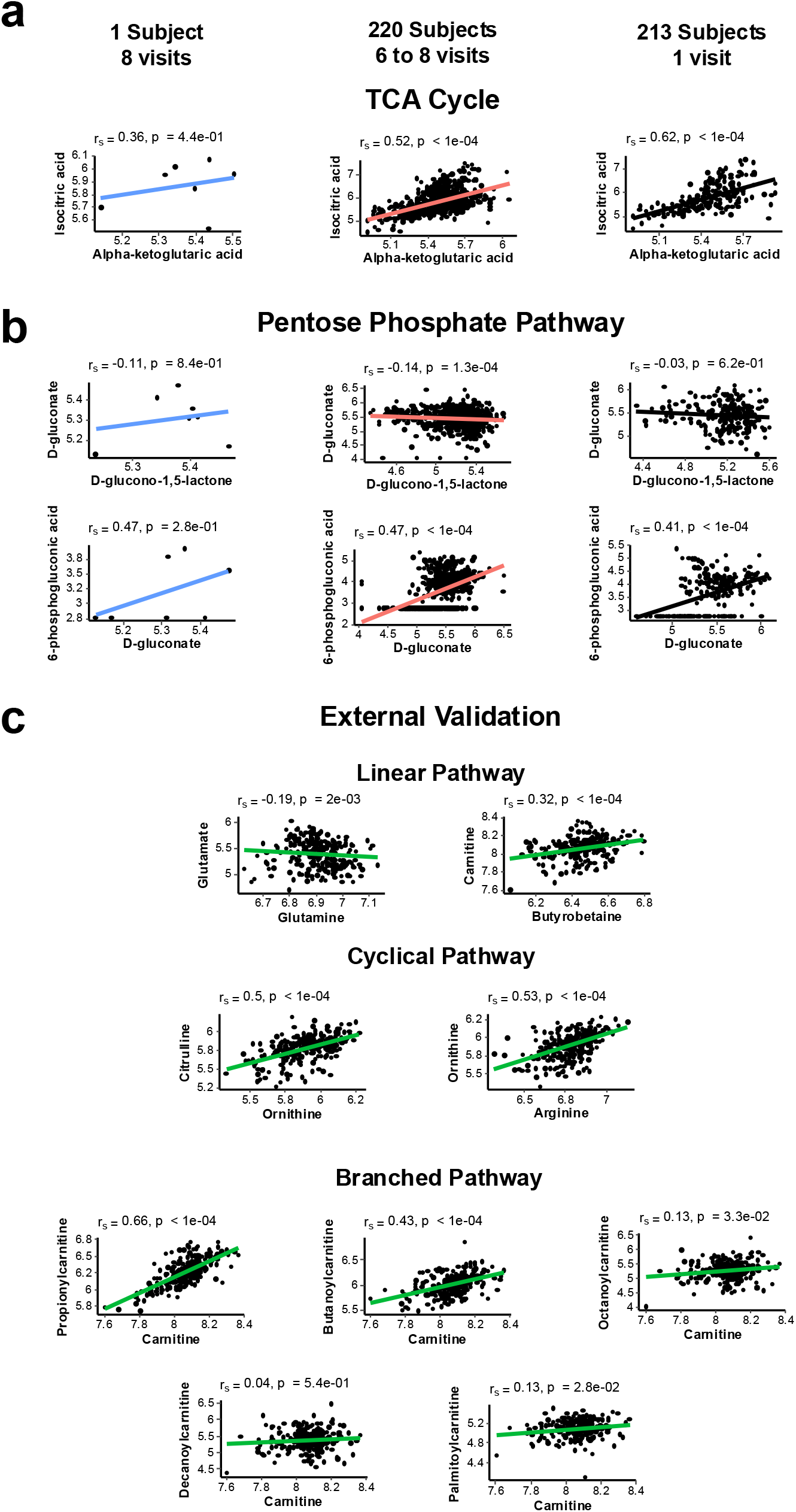
Additional precursor-product correlation tests in human plasma and serum. (**a**) Precursor-product correlation analyses in the TCA cycle pathway. Isocitric acid and alpha-ketoglutaric acid were tested for correlation. Data included eight repeated blood samplings from an individual collected over five years (blue, left), 220 individuals with multiple repeated blood samplings across five years (red, middle), and cross-sectional analysis of baseline samples for the 213 individuals in Panel **c** (black, right). (**b**) Precursor-product correlation analyses in the pentose phosphate pathway for D-gluconate and D-glucono-1,5-lactone (upper panel) and for 6-phosphoglyconic acid and D-gluconate (lower panels). Color schemes were the same as Figure 2a. (**c**) Precursor-product correlation analyses in multiple pathways (linear, cyclical, and branched) in an external validation dataset. This dataset represents a longitudinal study (3 timepoints) of 271 serum samples from 91 subjects with metastatic renal cell carcinoma treated with nivolumab.

As an external validation, we analyzed published LC-MS human serum data from a study consisting of 271 serum samples from 91 subjects with metastatic renal cell carcinoma treated with nivolumab.^10^ Nine RT and MS/MS confirmed precursor-product pairs were selected, similar to the pairs in Figure 1. Consistent with Figure 1, many precursor-product pairs were significantly correlated, while the pairs of decanoylcarnitine and carnitine and glutamate and glutamine were not significantly correlated (**Fig. 2c**).

In pathway analysis with tools such as Mummichog^2^, metabolite annotations have many-to-many relationships that complicate their interpretation. To explore how precursor-product relationships may enhance the pathway analysis, we performed untargeted pathway analysis with Mummichog^2^ in MetaboAnalyst^3^ on participants with obesity in the CHDWB cohort (CHDWB obesity).^11^ Using one-way ANOVA analysis with the *limma* algorithm,^12^ 1659 features (raw p-value < 0.05) out of 13112 total features were used for pathway analysis (**Fig. 3a**). The significantly altered valine, leucine, and isoleucine metabolism was selected for additional analysis. A correlation heatmap of all features in the Mummichog algorithm revealed multiple blocks of highly correlated features, with one block related to valine and leucine/isoleucine (**Fig. 3c**). The relevant network graph suggested three most relevant features associated with the confirmed valine feature (*m/z* 118.0866, 65 s, [M+H]^+^) (**Fig. 3c)**. Among the two annotated isoleucine features, one (*m/z* 132.1022, 51 s, [M+H]^+^) showed a higher correlation (Spearman correlation = 0.66, p < 0.0001) with the confirmed valine feature (*m/z* 118.0866, 65 s, [M+H]^+^) compared to the other correlations (**Fig. 3d)**. Similarly, among the three annotated valine features, the confirmed valine feature (*m/z* 118.0866, 65 s, [M+H]^+^) showed a higher correlation with the confirmed isoleucine feature (*m/z* 132.1022, 51 s, [M+H]^+^) compared to the other correlations (**Fig. 3d)**. These metabolites do not have enzymatic precursor-product relationships but are exchanged between cells and plasma by common transport systems. This correlation approach can be used to supplement pathway enrichment-guided annotations for improved accuracy.

**Figure 3.**
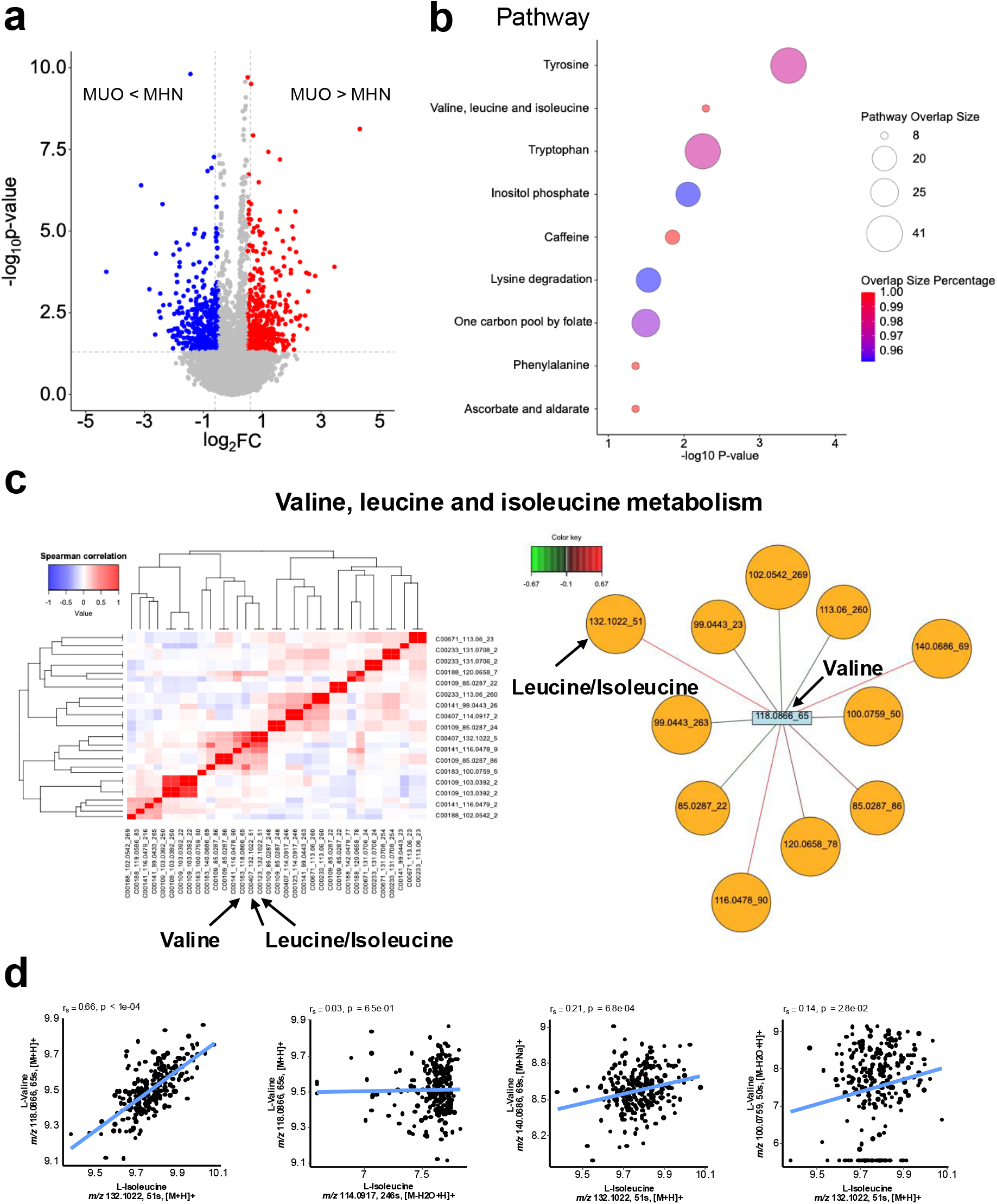
Untargeted pathway analysis in the CHDWB obesity study. **(a)** One-way ANOVA analysis with the *limma* algorithm was used to find significantly altered metabolomics features among subjects with metabolically healthy non-obesity (MHN), metabolically healthy obesity (MHO), and metabolically unhealthy obesity (MUO). Only the log2 fold changes between subjects with MHN and MUO were used in the volcano plot. A raw p-value of 0.05 was used to select significantly altered features. **(b)** Untargeted pathway analysis using the Mummichog algorithm. Untargeted pathway analysis was performed using the Mummichog version 1 (functional analysis without using retention time)^2^ in the MetaboAnalyst^3^. In the analysis, 1659 significantly altered features were used. **(c)** Correlation heatmap and network analysis among all valine, leucine, and isoleucine metabolism-related features in the Mummichog algorithm outputs. Spearman correlation was used in the correlation heatmap using the heatmap.2 function in the gplots R package (left). Partial Least Squares analysis in the mixOmics package was used between the confirmed valine feature and other features, and then a relevance network graph was drawn with a cutoff of 0.1 (right). **(d)** Precursor-product correlation analysis between annotated valine and annotated isoleucine features. Valine (*m/z* 118.0866, 65s, [M+H] ^+^) and isoleucine (*m/z* 132.1022, 51 s [M+H] ^+^) were confirmed by RT and MS/MS of the authentic standard.

## Discussion

As metabolomics moves beyond static pathway maps, enzymology-defined precursor-product correlations provide a mechanistically grounded view of metabolic pathways. Even though most metabolism occurs in organ systems other than blood,^1^ the plasma data (**Fig. 1, 2, 3**) showed that precursor-product relationships are preserved despite variable contributions of organ systems (**Fig. 1e**). Consistent precursor-product relationships on two in-house LC-MS datasets and an external metabolomics dataset show these correlations can be used across study designs, study populations, and various LC-MS methods.

Batch correction is often used to decrease the impact of analytical variation in population studies. We found that precursor-product correlations remained significant, although slightly attenuated, by the addition of batch correction by the ComBat algorithm (xMSanalyzer^13^) for the longitudinal and cross-sectional analyses of the 50 CHDWB subjects (**Supplementary Fig. 1**).

Given that the correlations in the study with 1 subject across 8 visits were mostly unchanged and all were analyzed in the same batch, it appears that ComBat batch correction may unintentionally disrupt biological covariations in the precursor-product relationships.

In comparison of glutamine and glutamate correlations among studies, we found that glutamine and glutamate were positively correlated CHDWB healthy plasma study, but not in the cancer serum study. This can occur because each of them is the precursor and product of many other metabolites, or it could occur because disease can distort the precursor-product correlation. For instance, cancer changes inter-organ glutamine trafficking by increasing the expression of glutaminases,^14,15^ which may explain the negative correlation result between glutamate and glutamine in the cancer study. A negative correlation between glutamate and glutamine also occurred in the CHDWB obesity study (data not shown), which may occur because glutamine is low in the subjects with obesity^16,17^. These findings suggest that precursor-product relationships in metabolomics analysis can have value for biological understanding even when precursors and products are not positively correlated.

In pathway enrichment analyses, such as those provided by Mummichog, one feature can represent many metabolites and complicate interpretation, even though the pathway-level results are reliable. Inclusion of precursor-product relationships provides a useful approach to improve reliability in the interpretation of the metabolite annotations. Importantly, the results with leucine, isoleucine, valine pathway metabolites show that metabolites which share cell transporter mechanisms can also display precursor-product like relationships even when they do not have enzymatic precursor-product relationships. This demonstrates the potential to extend the precursor-product relationship concept to metabolites sharing common transport systems, which was also described in another study.^18^

One must be cautious about enzyme-based relationships;^6^ some precursors and products do not show proportional behavior, and some distantly related metabolites have strong correlations.^19^ Thus, it may be arbitrary to connect two features together and claim they form a precursor-product relationship even if their chemical formula, chemical structures, and mass difference seem to support the claim. This is especially true for computational metabolomics, where data-driven metabolic network algorithms and in silico biochemical reactions were used for metabolite annotation. As demonstrated by **Figure 3c**, false positives will be mixed with true positives, and the number of annotation errors will increase via an iterative propagation process. From our perspective, known one-step precursor-product and transporter relationships should be prioritized compared to the data-driven metabolic network and in silico biochemical reaction approaches.

Overall, incorporating the precursor-product relationships into metabolomics analysis can improve pathway analysis and help metabolite identification. Extension to include metabolite transport systems and development of tools to improve capabilities will further realize the potential for comprehensive metabolomics to be a central hub for systems biology.

Metabolomics reflects all endogenous and xenobiotic alterations, genetic variations, enzyme activities, post-transcriptional regulation and protein modifications, diets, therapeutic drugs, diseases, and environmental exposures derived from biologic precursor-product relationships.^20–22^ The present results show that precursor-product relationships are highly preserved under different conditions and provide a valuable resource to enhance LC-MS-based metabolomics research.

## Resource availability

### Lead contact

Further information and requests for resources should be directed to the corresponding author, Dean P. Jones.

## Data and code availability

The datasets illustrated in this study were included in Supplementary Data 1 for Figure 1, Supplementary Data 2 for Figure 2, and Supplementary Data 3 for Figure 3. Raw data is available at the Metabolomics Workbench^23^: https://dev.metabolomicsworkbench.org:22222/data/DRCCMetadata.php?Mode=Study&StudyID=ST004055&Access=GecQ2393. The study is scheduled to be released on 2026-07-01 or before publication.

## Acknowledgments

This work was supported by the National Institute of Environmental Health Sciences (R01 ES031980 and P30 ES019776) and the National Institute on Aging (AG080247, AG085279, and U01 AG088658).

## Author contributions

G.S.M. established the study cohort. D.P.J. provided access to the metabolomics data. J.Z. analyzed the data. J.Z. and D.P.J. wrote the manuscript. Z.R.J., M.L.K., J.W., Y.M.G., G.S.M., and D.P.J. revised the manuscript. D.P.J. supervised all aspects of the project.

## Figure Legends

**Supplementary Figure 1. Precursor-product relationships in human plasma using ComBat batch corrected metabolomics data**. (**a**) Precursor-product correlation analyses among metabolites in multiple pathways for 8 blood collections for 1 subject across give years (blue). (**b**) Precursor-product correlation analyses among metabolites in multiple pathways among 50 subjects with repeated blood collections (6-8). **(c)** Precursor-product correlation analyses among metabolites in multiple pathways among 50 subjects with a baseline blood sampling. Data from 50 individuals with baseline blood collection (black) were used.

## STAR⍰Methods

### Study population

The metabolomics data are from the Center for Health Discovery and Wellbeing (CHDWB) study, a longitudinal cohort.^9^ CHDWB enrolled 668 participants who are essentially healthy employees at Emory University, and we selected 50 subjects with at least 6 visits in the first set of analyses (Fig. 1), and 213 subjects with only one visit for the second set of analyses (CHDWB obesity). Individuals with poorly controlled chronic disease or acute illness were excluded from the study enrollment. The first visit required overnight fasting for the blood draw, while the rest of the visits did not have the fasting requirement. The CHDWB study was reviewed and approved under Emory Investigational Review Board (IRB approval No. 00007243).

### Sample Preparation

50 μL of plasma sample was combined with 100 μL acetonitrile containing 11 stable-isotope internal standards added ([13C6]-D-glucose, [15N]-indole, [15N,13C5]-L-methionine, [2-15N]-L-lysine dihydrochloride, [15N]-choline chloride, [13C5]-L-glutamic acid, [13C7]-benzoic acid, [15N]-L-tyrosine, [15N2]-uracil, [3,3-13C2]-cystine, and [trimethyl-13C3]-caffeine). The mixture was kept on ice for 30 minutes, then spun at 14,000 × g for 10 minutes at 4 °C to precipitate proteins. The clear supernatant was transferred into autosampler vials and immediately loaded into a 4 °C autosampler for analysis.

### Metabolomics Instrumental Analysis

Metabolite profiling was performed using a Thermo Scientific Vanquish Duo HPLC system coupled to a Thermo Scientific ID-X high-resolution mass spectrometer operating in positive-mode electrospray ionization. Each sample was injected in triplicate, with pooled plasma references analyzed at the beginning, middle, and end of each sample batch to monitor analytical drift. Separation employed a Waters BEH Amide column (2.1 × 100 mm, 1.7 µm) with mobile phase A (0.1% formic acid, 1 mM ammonium acetate in water) and mobile phase B (95% acetonitrile, 0.1% formic acid, 1 mM ammonium acetate). The flow rate was 0.3 mL/min, starting at 90% B, linearly decreasing to 20% B between 0.6 and 3.15 min, then held for an additional 5 min. The mass spectrometer was set to 120k resolution, scanning from *m/z* 85–1275 on the Orbitrap detector. Sheath, auxiliary, and sweep gas flows were 50, 10, and 1 arbitrary units, respectively, with spray voltages of 3.5 kV. MS/MS spectra were acquired in a separate technical replicate using parallel reaction monitoring with targeted inclusion lists. Higher-energy collisional dissociation was performed at 35% normalized collision energy, collecting MS^1^ scans at 60k resolution and MS/MS scans at 30k resolution. For the CHDWB obesity study, in brief, the samples were analyzed by the HILIC chromatography paired with a Q Exactive HF Hybrid Quadrupole-Orbitrap mass spectrometer with positive ionization mode.^11^ For the cancer study used for external validation, in brief, the samples were analyzed with HILIC paired to a Q Exactive Orbitrap with positive ionization mode. Details were described elsewhere.^10^

### Metabolite Identification

Metabolites included in this study were confirmed by accurate mass MS^1^ signal (<u>±</u> 5 ppm), coelution with authentic standards (<u>±</u> 30 s), and ion dissociation spectra (MS/MS) matching authentic standards.^24^ These metabolites were the level 1 identifications according to the Metabolomics Standards Initiative and Schymanski criteria.^25,26^

### Statistical Analysis

Relative intensities of the metabolites were used for the correlation analyses. The intensities were median-summarized across triplicates, and non-detects were imputed using the half minimum method during this process. The intensities were log10 transformed and then used for the Spearman correlations between precursors and products. All precursor-product correlation analysis results were presented regardless of their correlation coefficients and p-values. A raw p-value of 0.05 was used in the Mummichog pathway analysis without p-value adjustment. A cutoff of 0.1 was used in the relevant network graph analyses.

