## Supplementary figures and images for "Enzymes define pathways and metabolic relationships"

### Supplemental Figure

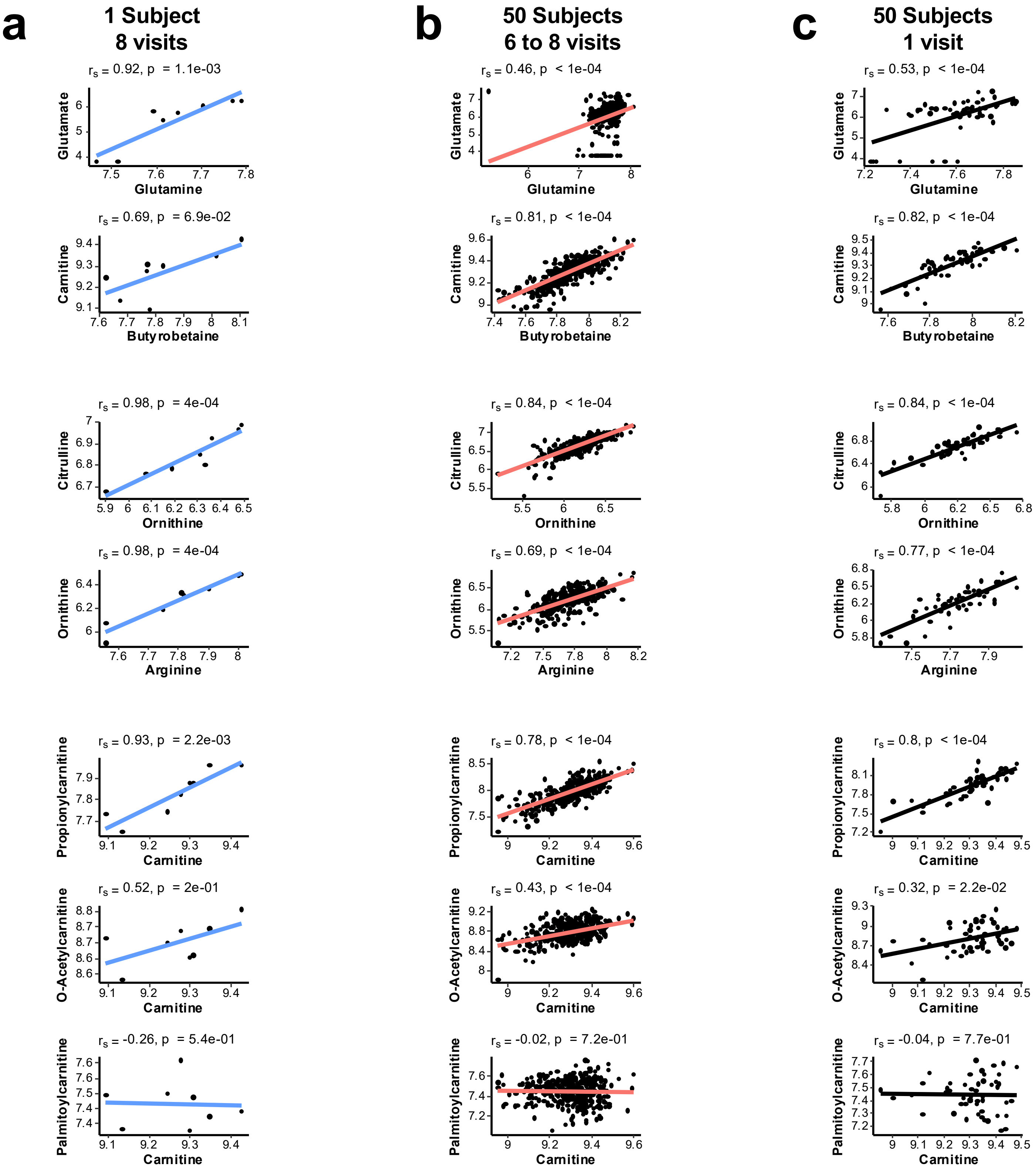
